# Human metapneumovirus SH protein promotes JAK1 degradation to impair host IL-6 signaling

**DOI:** 10.1101/2024.05.10.593594

**Authors:** Adam Brynes, Yu Zhang, John V. Williams

## Abstract

Human metapneumovirus (HMPV) is a leading cause of respiratory infections in children, older adults, and those with underlying conditions ^1,2,3,4^. HMPV must evade immune defenses to replicate successfully; however, the viral proteins used to accomplish this are poorly characterized. The HMPV small hydrophobic (SH) protein has been reported to inhibit signaling through type I and type II interferon (IFN) receptors *in vitro*, in part by preventing STAT1 phosphorylation^5^. HMPV infection also inhibits IL-6 signaling. However, the mechanisms by which SH inhibits signaling, and its involvement in IL-6 signaling inhibition are unknown. Here, we used transfection of SH expression plasmids and SH-deleted virus (ΔSH) to show that SH is the viral factor responsible for inhibition of IL-6 signaling during HMPV infection. Transfection of SH-expression vectors or infection with wildtype, but not ΔSH virus, blocked IL-6 mediated STAT3 activation. Further, JAK1 protein (but not RNA) was significantly reduced in cells infected with wildtype but not ΔSH virus. The SH-mediated reduction of JAK1 was partially restored by addition of proteasome inhibitors, suggesting proteasomal degradation of JAK1. Confocal microscopy indicated that infection relocalized JAK1 to viral replication factories. Co-immunoprecipitation showed that SH interacts with JAK1 and ubiquitin, further linking SH to proteasomal degradation machinery. These data indicate that SH inhibits IL-6 and IFN signaling in infected cells in part by promoting proteasomal degradation of JAK1 and that SH is necessary for IL-6 and IFN signaling inhibition in infection. These findings enhance our understanding of the immune evasion mechanisms of an important respiratory pathogen.

**Importance:** Human metapneumovirus (HMPV) is a common cause of severe respiratory illness, especially in children and older adults, in whom it is a leading cause of hospitalization. Prior research suggests that severe HMPV infection is driven by a strong immune response to the virus, and especially by inflammatory immune signals like interferons (IFN). HMPV produces a small hydrophobic protein (SH) that is known to block IFN signaling, but the mechanism by which it functions, and its ability to inhibit other important immune signals remains unexplored. This paper demonstrates that SH can inhibit another related immune signal, IL-6, and demonstrates that SH depletes JAKs, critical proteins involved in both IL-6 and IFN signaling. A robust understanding of how HMPV and related viruses interfere with immune signals important for disease could pave the way for future treatments aimed at mitigating severe infections.

## Introduction

Viruses invading a host are faced with physical, chemical, and immunological barriers to infection. Immunological barriers include interferons (IFN) and inflammatory modulators like IL-6 and TNF^6,7^. Each of these pathways is a complex multi-step process and thus viruses have evolved a wide variety of tools to interfere with signaling. Human metapneumovirus (HMPV) is an enveloped, single-stranded negative-sense RNA virus in the family *Pneumoviridae* discovered in 2001^8^. HMPV is a leading cause of acute lower respiratory infections in pediatric populations and the elderly with no approved treatments or vaccines^1,2,3,4^.

*Pneumoviridae* and the closely related *Paramyxoviridae* family inhibit several host innate immune pathways, with JAK/STAT signaling an especially common target. The V proteins of mumps virus, PIV5, and measles contribute to proteasomal degradation of signal transducer and activator of transcription (STAT) including STAT1 and STAT3 as a means of inhibiting IFN and IL-6^9,10,11^. The closely related respiratory syncytial virus (RSV) inhibits IFN production and signaling via its NS1 and NS2 proteins; however, these genes are absent in HMPV^12,13^. Instead, published data suggest the HMPV small hydrophobic (SH) protein inhibits type I and type II IFN signaling^14,15^. However, the mechanism of SH-IFN inhibition has not been described. Furthermore, HMPV infection was associated with reduced expression and increased proteasomal degradation of JAK1 and TYK2, factors important for both interferon and IL-6 signaling^16^. It has been reported that HMPV *infection* impaired IL-6-mediated STAT3 activation in a respiratory epithelial cell line^17^.

SH proteins of *Paramyxoviridae* and *Pneumoviridae* including PIV5, RSV, and HMPV have been established to inhibit TNF signaling or linked NF-ΚB activity activity^18,19,20^; however, there is little insight into the mechanisms by which this inhibition takes place. An exception to this is mumps virus SH, which inhibits TNF, TLR3, and IL-1β-mediated NF-KB activation by interacting directly with components of each of those receptor signaling complexes (TNFR1, RIP1, and IRAK1)^21^. Interestingly, mumps SH also interacts with the host ubiquilin4 protein (also known as A1UP), a factor implicated in facilitating proteasomal degradation of proteins^22,23^. This may suggest a broader mechanism by which other SHs, including that of HMPV, function. HMPV lacks accessorry proteins known to inhibit JAK/STAT signaling in related viruses. Given this, and the established ability of HMPV SH to inhibit interferon signaling, we hypothesize that SH is also responsible for IL-6 signaling inhibition. Additionally, we hypothesize that SH inhibits innate immune signaling by promoting proteasomal degradation of JAKs.

## Results

Stimulation of IL-6 or IFN signaling receptors leads to recruitment and activation of cellular JAKs (in the case of IL-6: JAK1, JAK2, and TYK2) which recruit and phosphorylate cellular STATs that form homo- or hetero-oligomers with transcriptional activator abilities. To assess whether HMPV SH could inhibit both STAT1 and STAT3 activation, we transfected IL-6 and type I IFN signaling-competent cells with vectors expressing either c-terminally myc-his tagged HMPV SH or a GFP control and performed short-term stimulation with either cytokine (**Fig 1**.). Expression of SH was confirmed by western blot using tag-specific antibodies (**Fig. S1 and Fig. 5** B.). Transient transfection and expression of HMPV SH, but not GFP, reduced stimulated phosphorylation of STAT1 and STAT3. We measured the IFN-stimulated gene (ISG) *Mx1* by qPCR and found that IFN-induced Mx1 mRNA was reduced by transfection with SH but not GFP (**Fig. S2**). This confirmed the ability of SH to inhibit type I IFN-mediated STAT1 activation and demonstrated that ectopically expressed SH impaired IL-6 signaling *in vitro*.

**Figure 1.**
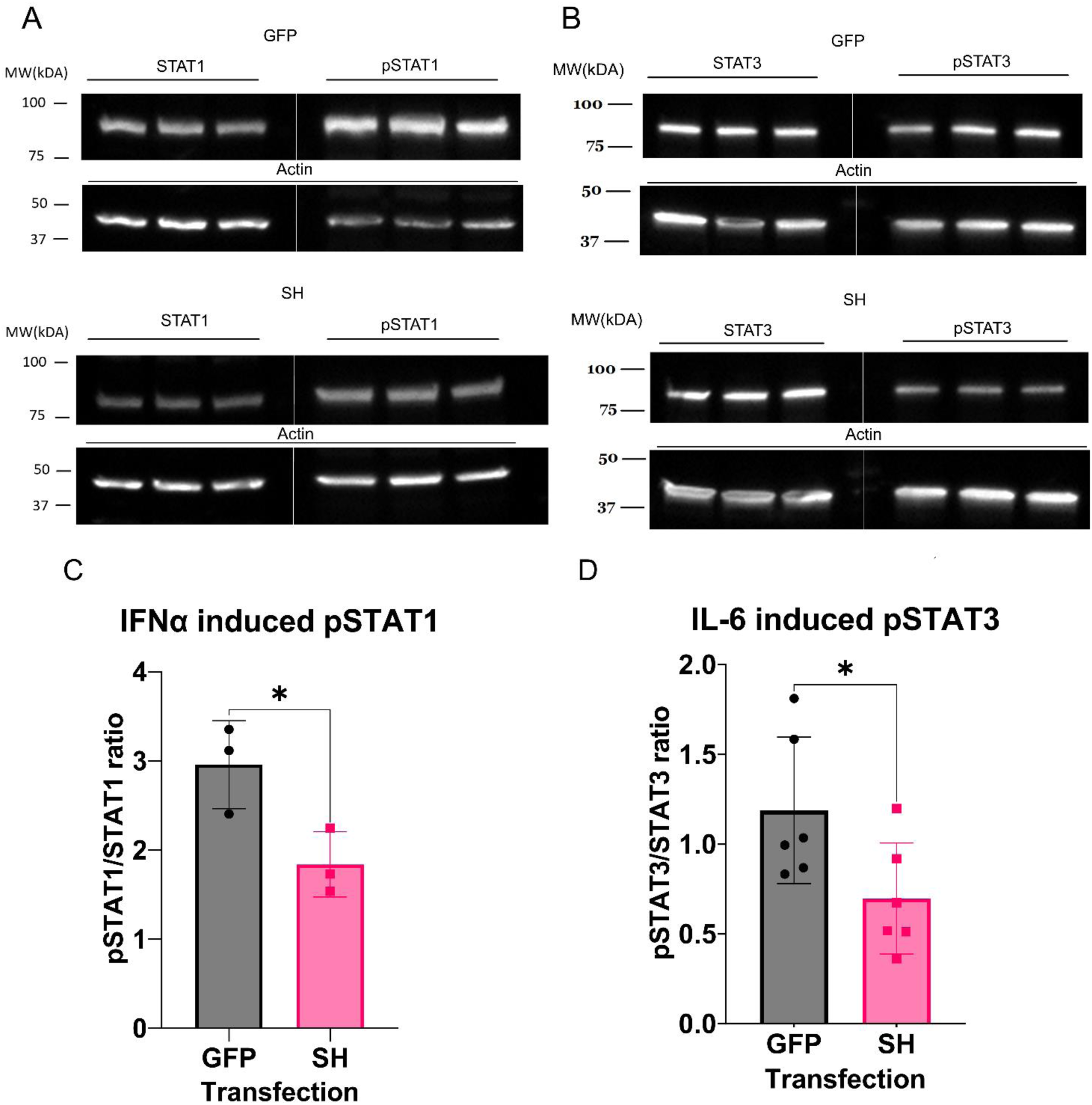
HMPV SH expression impairs IFN and IL-6 signaling. 293T cells were transfected in triplicate with pcDNA3.1+ encoding either c-terminally myc-his tagged SH or GFP as labeled. Cells were treated 30 hours post-transfection with either 1000 units/mL human IFN-αa2 (A & C) or 20ng/mL human IL-6 (B & D) for 20 minutes. Total cellular lysate was collected and used to generate western blots probing for total STAT1 or activated pSTAT1 Y701 (A) or total STAT3 or activated pSTAT3 Y705 (B). The ratio of phosphorylated STAT to total STAT was calculated for each replicate and the results of all biological replicates performed are shown in C & D. Statistical analysis was performed with Student’s t-test on six biological replicates. Error bars represent SEM. * = P<0.05.

We next sought to demonstrate whether SH expression was required to inhibit IL-6 signaling during viral infection. A respiratory epithelial cell line was infected with wildtype (WT) HMPV 94-49 or one of two recombinant HMPV strains: rWT94-49 expressing an otherwise wildtype C-terminally HA-tagged SH gene, or ΔSH 94-49 generated by introducing three stop codons near the beginning of the SH gene. Recombinant WT and ΔSH 94-49 replicated with comparable efficiency to the parental WT 94-49 and produced or failed to produce tagged SH during infection as expected (**Fig. S3**).

Respiratory epithelial cells were infected with WT HMPV, recombinant WT HMPV, ΔSH HMPV, or mock cell lysate, and treated with IL-6 or a PBS (vehicle) control. IL-6-mediated STAT3 activation was significantly reduced relative to mock infection in WT and recombinant WT-infected cells but not ΔSH-infected cells as assessed by western blot (Fig. 2). Combined with the transient transfection experiments, this indicates that SH is both necessary and sufficient for IL-6 signaling inhibition.

**Figure 2.**
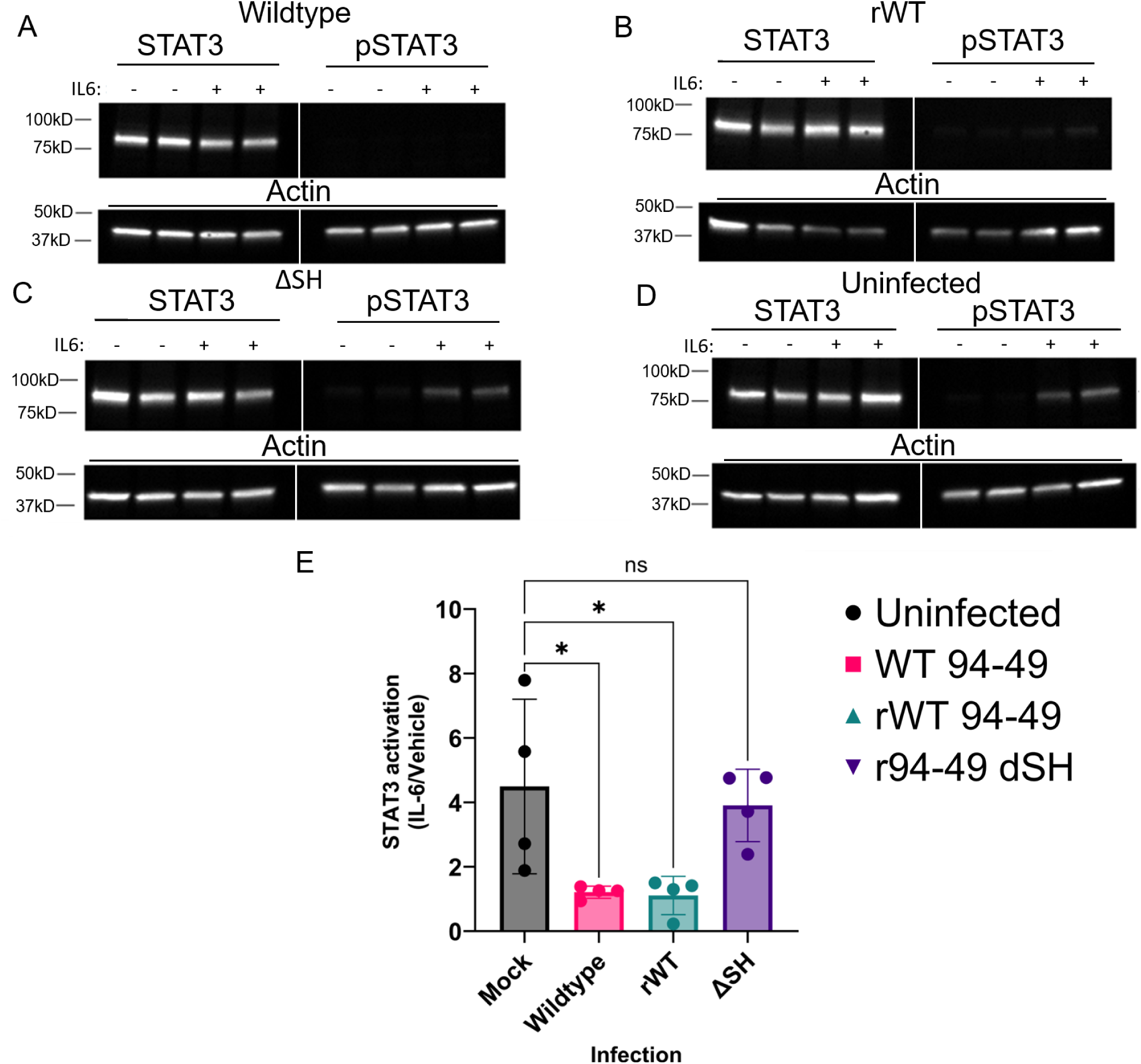
SH expression during infection is essential for IL-6 signaling inhibition. A549 cells were infected at an MOI of 0.5 for 24h with one of the four listed HMPV 94-49 strains: parental WT (A), recombinant WT (rWT) consisting of recombinant 94-49 with a C-terminal HA tag on the SH gene (B), ΔSH consisting of rWT with three stop codons inserted near the start of the SH gene (C), or uninfected (D). At 24hpi, cells were treated for 20 minutes with 20ng/mL human IL-6 and harvested. Lysate was blotted for total STAT3 and activated phospho-STAT3. The ratio of STAT3 activation (phospho-STAT3 to total STAT3 signal) in IL-6 to mock cytokine (PBS)-treated cells is quantified in E as “STAT3 activation (IL-6/Vehicle)”. Statistical analysis represents ordinary one-way ANOVA performed on four biological replicates. Error bars represent SEM. * = P<0.05.

Mumps virus and other paramyxoviruses promote proteasomal degradation of host STATs using V proteins^24,25^. While HMPV lacks an identified V protein, viral infection reduced levels of JAK1 and TYK2 protein via proteasomal degradation^16^. To investigate whether SH was required for reduced expression of JAKs, A549 cells were infected with WT or ΔSH mutant virus, and levels of JAK1 protein were measured by Western blot (Fig. 3). There was a significant decrease in JAK1 levels in either SH-expressing infection (rWT or WT) compared to ΔSH infection, but not in SH-transfected cells. Levels of JAK1, JAK2, and TYK2 were also assessed in HMPV vs mock-infected A549 cells, with a significant infection-mediated reduction noted in JAK1 and TYK2 but not JAK2 (**Fig. S4**). We measured JAK1 RNA by RT-qPCR and found an increase in JAK1 RNA in SH-expressing infections (**Fig. S5**), in contrast to the reduction in JAK1 protein in the presence of SH protein. Taken together, these data suggest an SH-dependent increase in JAK1 turnover in infected cells.

**Figure 3.**
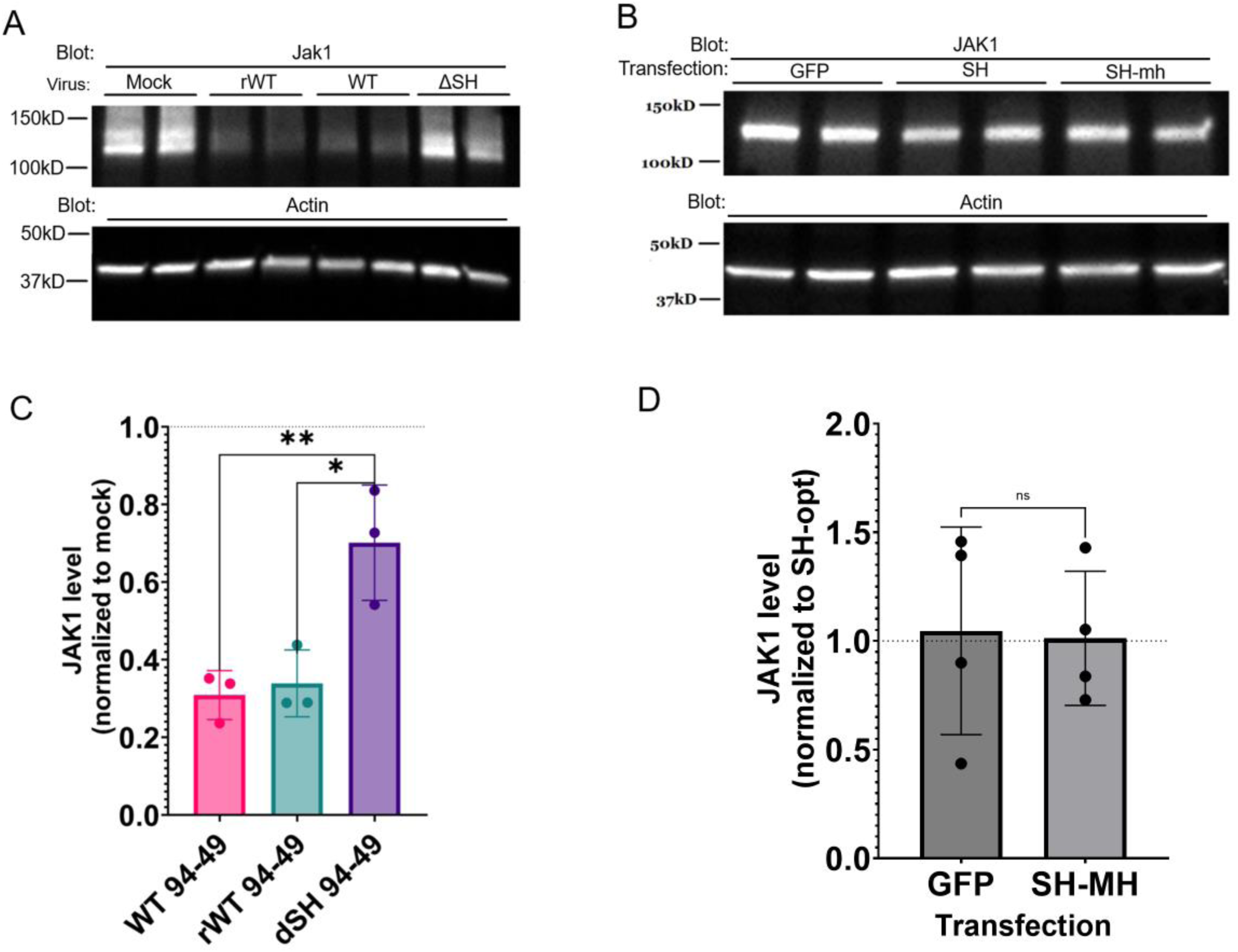
SH expression reduces JAK1 levels in HMPV infection but not SH transfection. A549 cells were infected at an MOI of 0.5 for 24 hours & JAK1 levels were assessed by western blot (A & C). 293T cells were transfected with vectors encoding a GFP transfection control, codon-optimized 94-49 HMPV SH, or codon-optimized, C-terminally myc-his tagged 94-49 HMPV SH. JAK1 levels were normalized to untagged-codon optimize SH-transfected conditions (SH-opt) and assessed by western blot at 30 hours post-transfection (B & D). Statistical analysis represents a one-way ANOVA performed on three (C) biological replicates or a t-test performed on four biological replicates (D). Error bar represents SEM. * = P<0.05, **= P<0.01.

Therefore, we assessed whether the SH-mediated decrease in JAK1 protein could be attributed to proteasomal degradation. A549 cells were infected with WT, recombinant WT, or ΔSH virus, and treated with either lactacystin (a specific, irreversible proteasome inhibitor^26^) or DMSO vehicle control. The ratio of JAK1 levels in proteasome inhibitor-treated vs. mock-treated conditions was used to measure the level of JAK1 proteasomal degradation in each infection condition. Proteasome inhibition led to an almost 4-fold increase in JAK1 levels after WT infection (Fig. 4) indicative of proteasomal degradation of JAK1 during HMPV infection. Significantly less proteasomal targeting of JAK1 was noted in ΔSH infection relative to WT; however, the difference between recombinant WT and ΔSH infection did not reach the level of significance. The higher level of JAK1 proteasomal degradation in WT and recombinant WT-infected cells is consistent with SH promoting proteasomal degradation of JAK1 protein.

**Figure 4.**
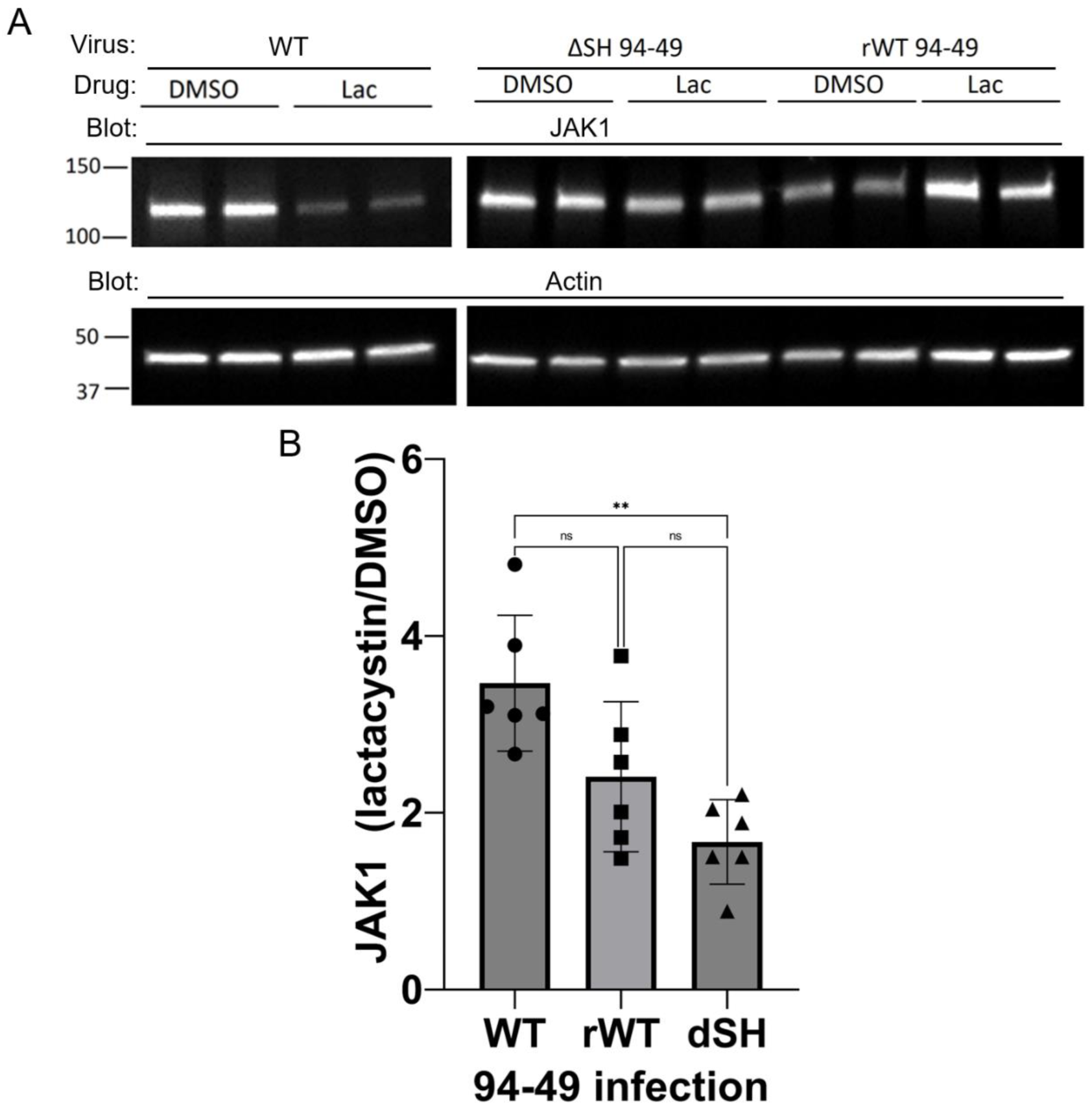
SH may promote proteasomal degradation of JAK1. A549 cells were infected with indicated HMPV 94-49 strains at MOI = 0.5 with proteasome inhibitor lactacystin (Lac) or mock treatment added 1h post-infection. Cells were harvested at 18h post-infection and JAK1 levels were assessed by western blot. The ratios of JAK1 measured in proteasome inhibitor-(Lac) treated vs mock (DMSO) conditions are shown as a measure of proteasomal degradation in each infection condition (B).

We then sought to determine whether SH associated directly with JAK1, as well as components of the ubiquitin-proteasome pathway. 293T cells were transfected to express untagged HMPV SH, C-terminally myc-his tagged SH, or GFP control. SH and JAK1 were separately precipitated using protein or tag-specific antibodies, and eluted protein was analyzed by immunoblot. This process was repeated for ubiquitin, with cells transfected to express tagged SH or GFP control followed by SH or ubiquitin precipitation (Fig. 5). High molecular weight species of SH, previously published as SHg2^27^ and thought to represent highly glycosylated species of the protein, precipitated with ubiquitin and JAK1. Given that bands identical in molecular weight stained positive for ubiquitin after SH precipitation, this suggests that these high molecular weight SH species represent ubiquitinated SH in addition to or instead of glycosylated protein. This polyubiquitination may enable further interactions between SH and components of the ubiquitin proteasome system, facilitating degradation of host factors.

**Figure 5.**
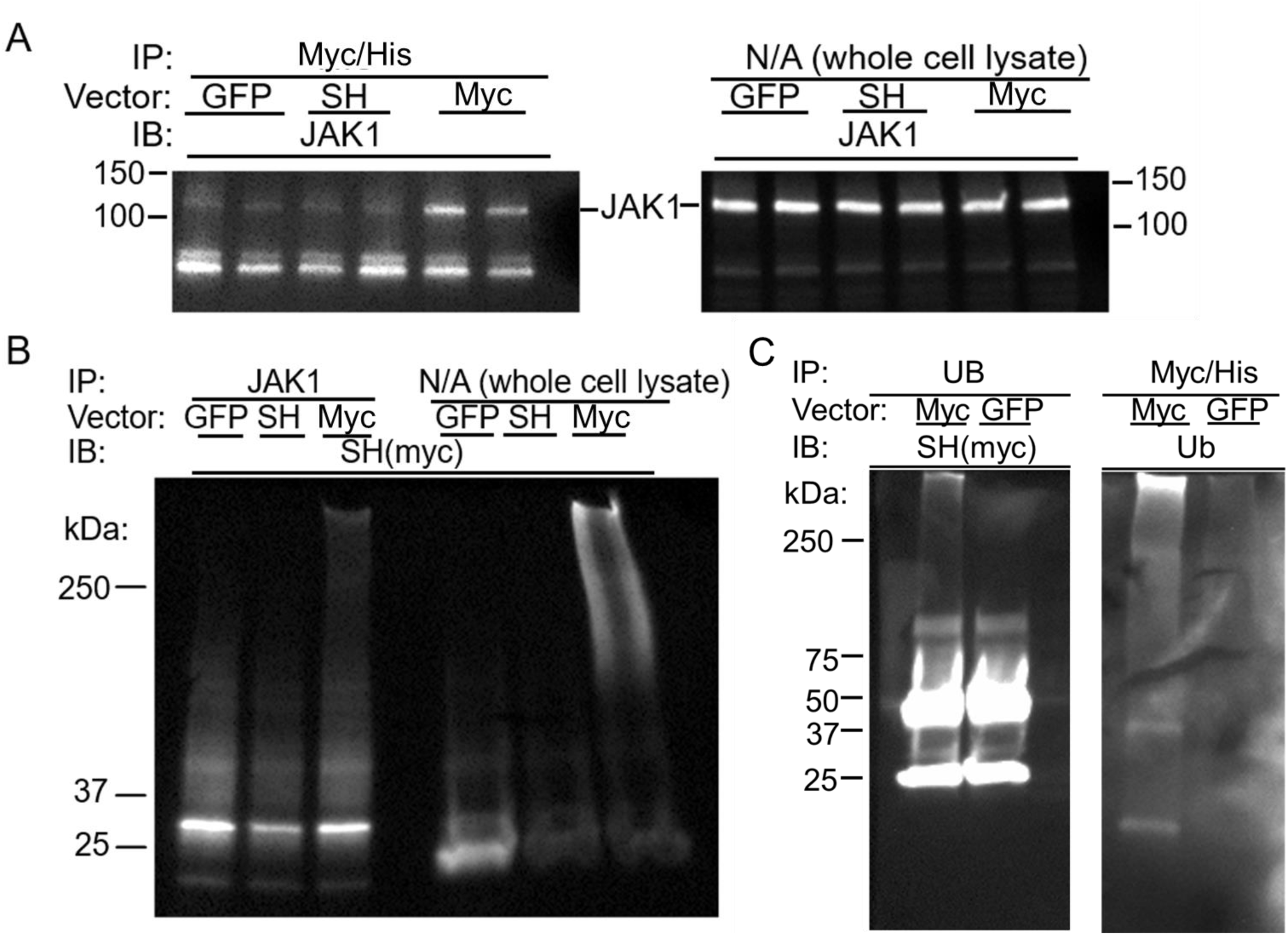
HMPV SH co-precipitates with JAK1 and ubiquitin. 293T cells were transfected to express GFP, SH, or C-terminally myc-his tagged SH (myc) as indicated. SH (A), JAK1 (B), or Ubiquitin/Ub (C) were precipitated using tag-, JAK1-, or ubiquitin-specific antibodies respectively. Eluted proteins were analyzed by western blot. “IP” indicates protein/tag precipitated, “IB” indicates the protein/tag blotted for.

We sought to further characterize SH-JAK1 interactions using confocal microscopy of infected cells. JAK1 and HMPV N (a marker of infection and a subcellular marker of viral inclusion bodies^28^) were localized within HMPV or mock-infected respiratory epithelial (A549) cells. As expected, punctate and globular inclusions staining positive for HMPV N were detected in all infected cells (Fig. 6).

**Figure 6.**
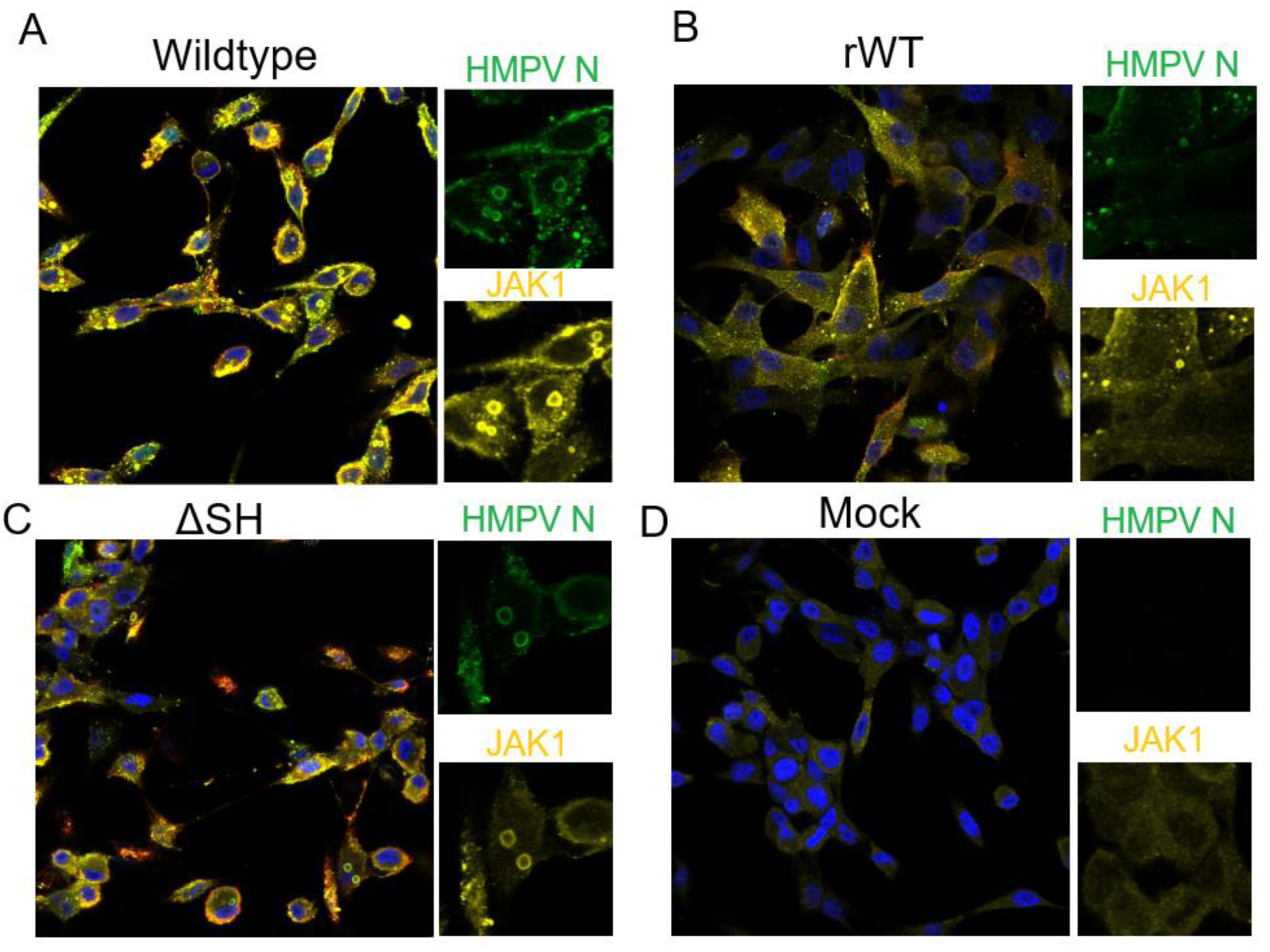
JAK1 localizes to viral inclusion bodies during infection. A549 cells were infected for 24 hours with an MOI of 0.5 of parental WT HMPV (A), recombinant WT (B), ΔSH (C), or mock virus (D), and stained for JAK1, and HMPV N.

These inclusions, known as viral inclusion bodies (IB) or replication factories are produced ubiquitously in infections by Mononegavirales, including *Paramyxoviridae* and HMPV^28,29,30,31,32^. IBs are liquid-liquid phase-separated structures thought to assist in viral genome replication or particle assembly, but have been implicated in host immune evasion; RSV sequesters dsRNA sensing proteins in cytoplasmic IB with subsequent destruction facilitated by NS1 and NS2 proteins^33,12^. We therefore decided to probe whether HMPV SH might prevent JAK1 activity via a similar mechanism.

Microscopy revealed that following infection with all HMPV strains (parental WT, recombinant WT, or ΔSH) JAK1 relocalized from diffuse cytoplasmic staining in mock infection to punctate structures colocalized with N-stained IB. When infected cells were stained with JAK1 antibody alone, JAK1 foci were retained. Additionally, staining with a separate monoclonal JAK1 antibody still revealed infection-dependent JAK1 puncta (**Fig. S6**). Fluorescence bleed-through from the HMPV N channel to the JAK1 channel was further refuted by imaging the same field of infected cells with the HMPV N (Alexafluor 488) laser activated or deactivated, confirming that JAK1 signal was independent of HMPV N signal. Furthermore, partial knockdown of JAK1 with siRNA attenuated JAK1 fluorescence relative to a non-targeting SI control (data not shown).

## Discussion

HMPV is a ubiquitous respiratory pathogen responsible for significant morbidity in pediatric, immunocompromised, and older adult populations^1,2,3,4,34^. Immune and inflammatory pathways are critical to HMPV disease^35,36^. Prior research indicated that HMPV infection could inhibit several host immune signaling pathways. HMPV G protein expression inhibits RIG-I-mediated IFN production^37^ while the M2-2 protein inhibits MAVS, a host factor critical for viral RNA recognition^38^. HMPV SH protein inhibits TNF-mediated NF-ΚB activation, as well as type I and II IFN-mediated STAT1 activation^5,39^. HMPV infection promoted proteasomal degradation of JAK1 and TYK2, though a mechanism was not described^16^. Given that HMPV infection has been implicated in inhibition of another JAK-STAT signaling pathway, IL-6, a shared mechanism involving SH is plausible.

This research confirmed SH’s ability to inhibit IFN signals and established a role for SH in inhibiting IL-6 signaling. SH was both necessary (in infection) and sufficient (in transfection conditions) to inhibit IL-6 signaling. However, the magnitude of the reduction in STAT3 activation in SH vs. GFP transfected cells (∼2-fold) was markedly smaller than that in wildtype infected vs ΔSH infected cells. While the presence of untransfected cells, and differences between the A549 and 293T cells used in these experiments may account for some of this difference, some may be due to SH cooperating with other viral proteins or infection processes to exercise full inhibitory activity. This is indicated by the ability of wildtype infection, but not infection with ΔSH virus or transfection with SH alone, to decrease cellular JAK1. Our data confirm degradation of JAK1 and TYK2 by HMPV infection.

Moreover, we found a significant SH-dependent reduction in JAK1, with a trend towards significant SH-mediated proteasomal degradation of JAK1. The additional observation that SH co-precipitated with JAK1 itself as well as ubiquitin, a component of the ubiquitin-proteasome system further suggests that SH may adapt JAKs for degradation.

Confocal microscopy begins to indicate how, when, and where SH may be promoting proteasomal degradation of JAK1. Many RNA viruses form phase-separated inclusion bodies thought to facilitate genome replication^,30,31,32^. In RSV, a close HMPV relative, immune sensing factors RIG-I and MDA5 were relocalized to viral inclusion bodies^33^. We show that HMPV possesses a similar ability, with infection relocalizing JAK1 to viral inclusions. While this relocalization was not SH-dependent, it is consistent with SH working in concert with other infection processes to impair host immune signaling. During infection, other viral proteins may recruit JAK1 to viral replication factories, where SH adapts the protein to the ubiquitin-proteasome pathway, resulting in its destruction. As infection without SH is insufficient to block JAK1-dependent signaling, despite localization of JAK1 to IB, relocalization of JAK1 during infection may be incomplete. There may be an equilibrium between IB-localized and non-localized JAK1, with the residual non-localized JAK1 being sufficient to participate in signaling in infected cells. By promoting degradation of JAK1, possibly at the IB, SH may shift this equilibrium in favor of more complete IB-localization of JAK1, interrupting signaling. Alternately, it may serve to mop up residual non-IB localized JAK1 to much the same effect. It is also possible that SH may enhance JAK localization to inclusion bodies.

Given the significant public health toll of HMPV and the link between IFN production and severe HMPV disease, understanding how this pathogen evades host defenses and IFN signaling is important^35^. We have established here that HMPV SH is the viral factor responsible for IL-6 signaling inhibition and that SH expression in infection promotes a likely proteasome-mediated depletion of cellular JAK1 and TYK2. In concert with this, infection relocalizes JAK1 to viral inclusion bodies which may serve as a site for degradation or sequestration of JAKs. Taken together, these results enhance our understanding of immune inhibition by a prevalent respiratory pathogen of public health significance.

## Materials and Methods

### Cell lines and virus

A549 cells (ATCC^®^ CCL-185™), LLC-MK2 (ATCC^®^ CCL-7™) cells, and 293T cells (ATCC^®^ CRL-3216™) were obtained from ATCC. Human cell lines were confirmed by STR genotyping. A549 cells and 293T cells were maintained in DMEM (Corning) supplemented with 10% FBS. LLC-MK2 cells were maintained in Opti-MEM (Gibco) supplemented with 2% FBS. Viruses including WT HMPV (clinical strain TN/94-49, A2 genetic lineage) were grown and titered in LLC-MK2 cells as described^40^. To generate recombinant 94-49, viral genomic plasmid was generated as previously described^41^. A 3’ double HA tag sequence and/or 3 stop codons were introduced to the SH gene (for ΔSH and rWT respectively) by PCR primer-directed mutagenesis. Modified SH gene was integrated into the genomic plasmid via HiFi assembly as per manufacturer recommendations [NEB, E5520S]. Cells were infected by removing media and incubating with virus diluted in Opti-MEM containing 0% serum, 5ug/mL trypsin, and 100ug/mL CaCl2. All media was supplemented with l-glutamine, gentamicin, and amphotericin.

### Plasmids and transfection

Plasmids used were generated as described^5^ with the exception that the SH used in these experiments was C-terminally tagged with myc-his dual tag rather than GFP or left untagged as indicated. GFP transfection controls used the pcDNA3.1 + C-terminal GFP tag plasmid without a gene insertion.

Plasmid transfection took place in 12-well plates. Cells to be transfected were grown to 70-90% confluence and transfected with the indicated plasmid as per the Transit-LT1 (Mirus bio, MIR 2304) manufacturer recommendations with the exception that 2ug DNA and 6uL transfection reagent were used per well.

### Western blot

Cells to be used for western blot were washed in PBS and lysed in ice-cold RIPA lysis buffer. Lysates were sonicated to disrupt genomic DNA, and protein content was analyzed via DC protein assay (Bio Rad, 5000111). Lysates were prepared for blotting by diluting 30ug lysate protein in water and reducing Laemmli sample buffer (Boston Bioproducts, BP-111R). Ladder samples were prepared using 8uL/lane ladder (BioRad, 1610373). Protein samples were heated to 95°C for 3 minutes, cooled to room temperature, and run on 4-20% SDS-PAGE gradient gels (BioRad, 4561094). Gels were transferred to PVDF membranes & blocked for 1h in TBST + 5% nonfat powdered milk. Blocking was followed by primary and secondary antibody incubation (rocking overnight at 4°C and 1h at room temperature, respectively) with TBST washes following each antibody incubation. Antibodies used were JAK1 (Cell Signaling Technology, 50996S), STAT1 (Cell Signaling Technology, 14994S), pSTAT1 Y701 (Abcam, Ab109457), STAT3 (Cell Signaling Technology, 12640S), pSTAT3 Y705 (Cell Signaling Technology, 9145S), myc (CST, 2276S), β-actin (CST, 3700S) anti-rabbit (CST, 7074S), and anti-mouse (CST, 7076S).

### Immunoprecipitation

In cells transfected to express SH or a control protein, precipitation was accomplished using either non-denaturing lysis buffer or reversible cross-linking of cellular proteins followed by lysis with denaturing lysis buffer (RIPA). In crosslinking IP, transfected cells were crosslinked with DSP (Millipore Sigma, D3669-500MG) as per manufacturer recommendations from ProteoChem. Briefly, cells were washed in PBS and crosslinked in 2mM DSP for 30 minutes at room temperature. The reaction was quenched by the addition of 50mM tris pH 7.4 and cells were lysed and immunoprecipitated as per Abcam immunoprecipitation protocol. For non-denaturing IP, crosslinking was omitted before following the IP protocol, and the indicated non-denaturing lysis buffer was used for lysis. Pre-clearing was accomplished using an excess of host-isotype antibody-conjugated Sepharose beads (Cell Signaling Technology, 3420S & 3423S) at a 1:20 concentration. SH was precipitated using Sepharose beads conjugated to anti-myc antibody (Cell Signaling Technology, 3400s). Jak1 and ubiquitin IP antibodies were (Cell signaling Technology, 50996s; and eBioscience, 14-6078-82) respectively. IP antibodies were used at 1:200, 5ug/mg lysate, and 1:200 respectively for the above antibodies. Antibodies were conjugated in lysate to protein G-coupled agarose beads (Cell Signaling Technology, 37478P). Glycine buffer elution was used following IP as per the linked protocol and eluate was processed as for western blot (note that the sample buffer used for blotting contains β-mercaptoethanol, reversing DSP crosslinking).

### Confocal microscopy

Cells were fixed in 4% formaldehyde for 10 minutes and permeabilized in PBS + 0.2% Tween-20 [Sigma Aldrich, P1379] for 10 minutes. Permeabilized cells were incubated with primary antibody for 1h followed by washing in PBS. Cells were incubated in fluorescently conjugated secondary antibodies for 30 minutes, stained with DAPI at 1ug/mL for one minute, washed, and mounted to a coverslip in Aqua Polymount (Polysciences, 18606-20). Imaging was accomplished on a Leica Stellaris 5 confocal microscope. Antibodies used were JAK1 (Thermo Fisher PA5-115442), monoclonal JAK1 (Millipore Sigma, 05-1154), HMPV N (EMD Millipore, MAB80138), HMPV F (54G10 monoclonal^42^ generated by Genscript).

### qPCR

RNA was extracted from total cellular lysate (Invitrogen, 12183025). qPCR was performed on an Applied Biosystems Quantstudio 5 machine using the AgPath-ID One-Step RT-PCR kit (Applied Biosystems, 4387391) and Taqman primers as per manufacturer recommendations. Thermo Fisher Taqman primers were used against HPRT1 (Hs02800695_m1), JAK1 (Hs01026983_m1), TYK2 (Hs01105947_g1), and MX1 (Hs00895608). RNA fold change (RQ) was calculated for sample groups in the Applied Biosystem Quantstudio Design and Analysis software. HPRT1 was measured as a housekeeping gene unless otherwise noted.

## Acknowledgments

We would like to thank Drs. Neal Deluca, Rachel Gottschalk, Amy Hartman, Fred Homa, Larry Kane, and Saumen Sarkar for their advice, and Dr. Jorna Sojati for protocols. This work was supported by funding from NIH AI085062, the Henry L. Hillman Foundation (JVW), and NIH T32 AI-049820.

## Supplemental figures

**Figure S1.**
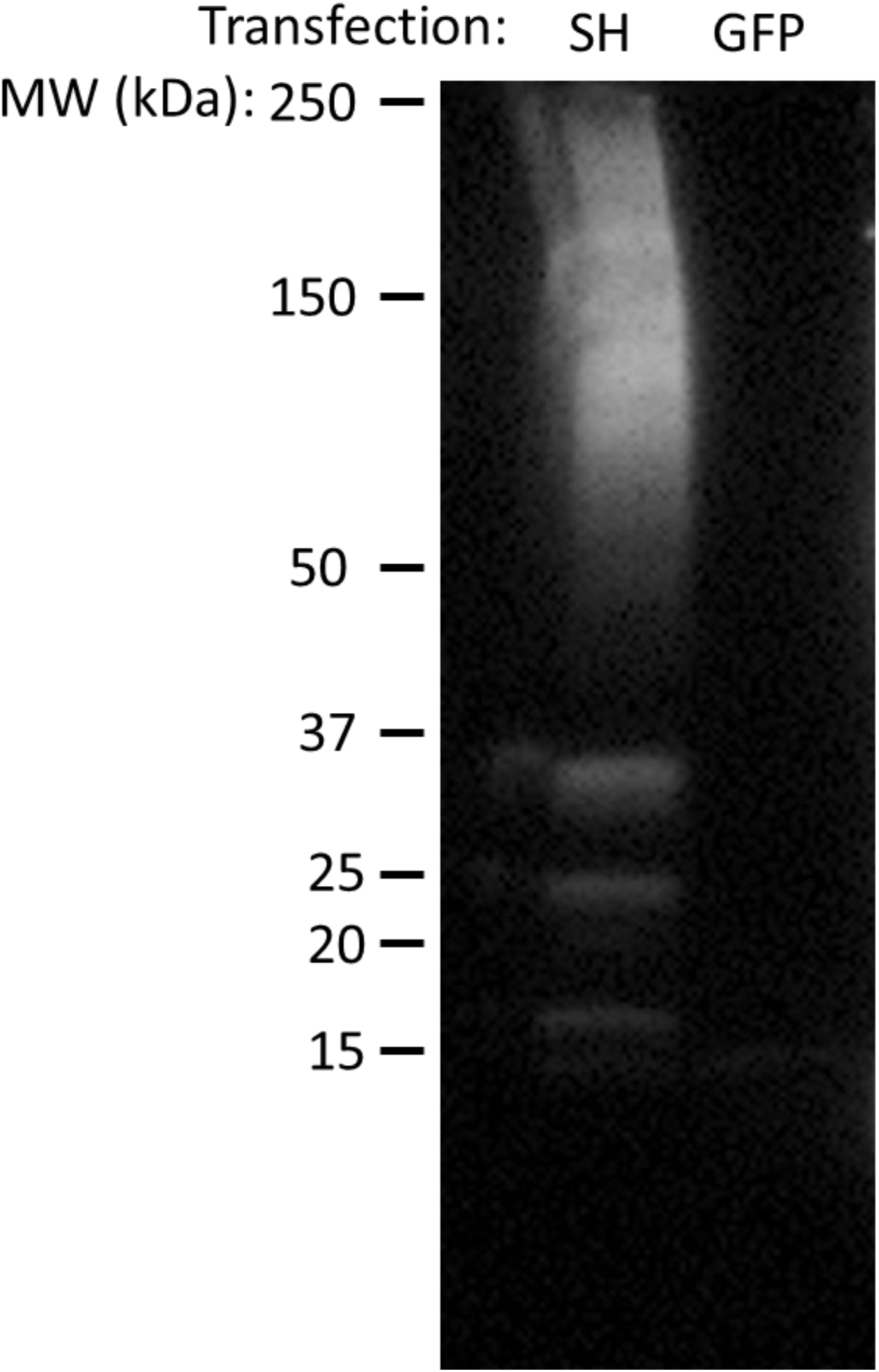
293T cells were transfected with pcDNA3.1+ encoding either c-terminally myc-his tagged SH (SH) or GFP as labeled. Cells were harvested 30 hours post-transfection and blotted for myc tag. Note the SH species of similar molecular weights observed in Biacchesi et al. 2004, JVI.

**Figure S2.**
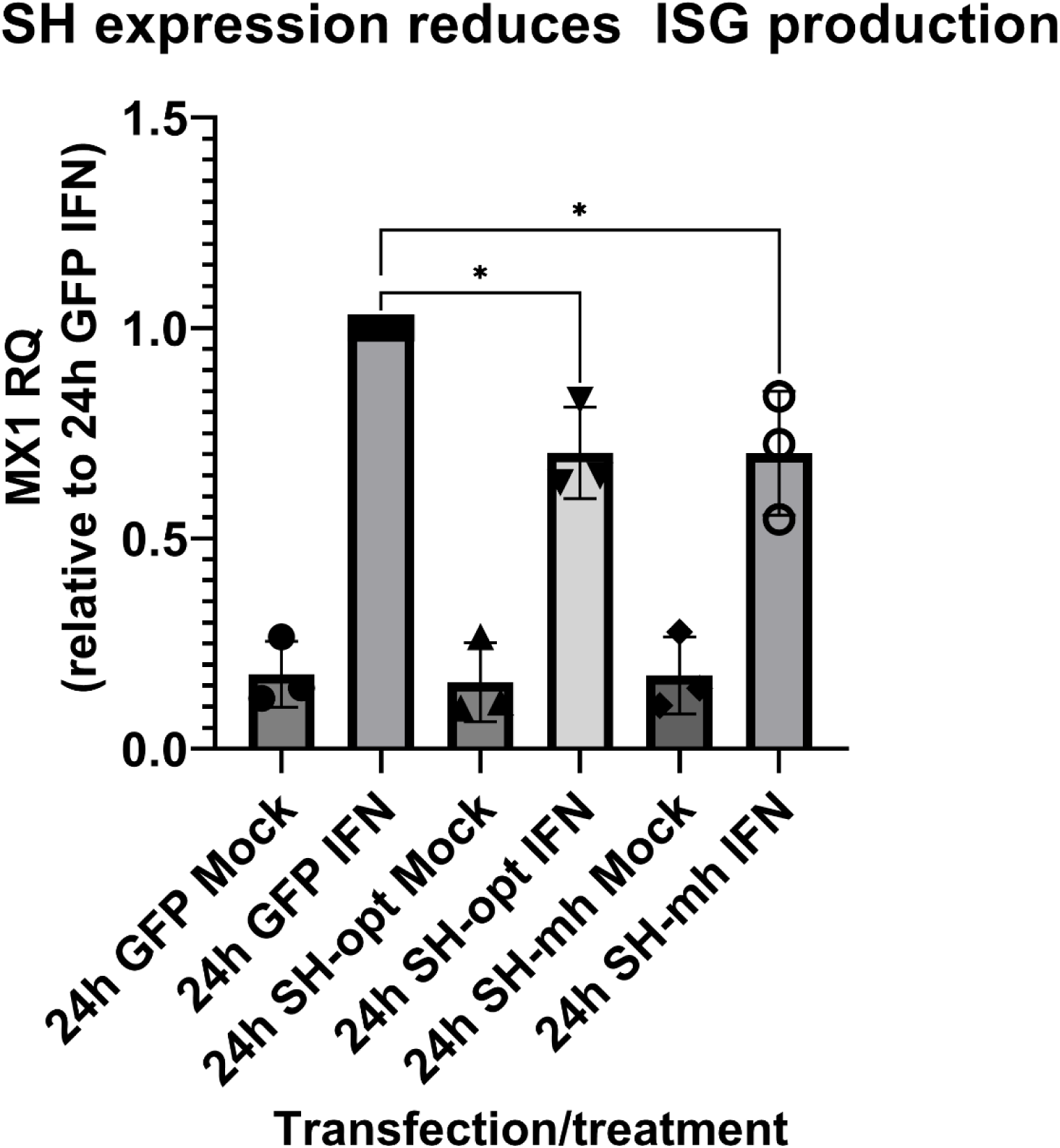
293T cells were transfected with either pcDNA3.1+ GFP or pcDNA3.1+ expressing either codon-optimized (opt) or codon-optimized, C-terminally myc-his tagged 94-49 SH. 30 hours post-transfection cells were treated for 0h or 24h with mock cytokine or 1E3units/ml IFN-αa2 (0h data not shown). MX1 ISG RNA was quantified by qPCR. RQ normalized to 24-hour GFP-transfected IFN treatment group. Bars represent N=3 with one-way ANOVA for statistical comparison.

**Figure S3.**
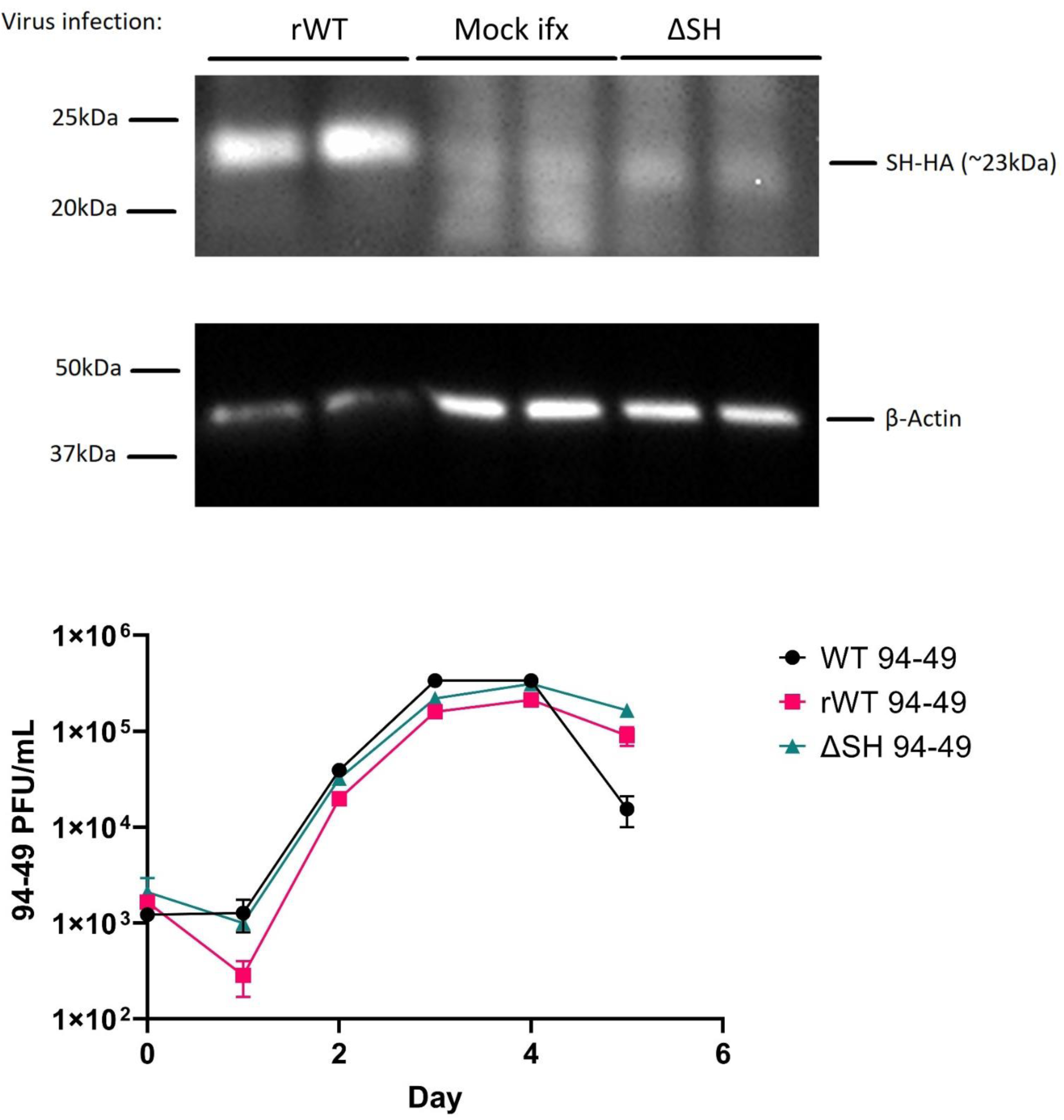
(A) LLC MK2 cells were infected at an MOI of 1 for 24h with two independently generated stocks of recombinant WT HMPV 94-49 (here referred to as 5 & 7), ΔSH 94-49, or mock. Lysate collected 24hpi was blotted for SH’s C-terminal HA tag. (B) LLC-MK2 cells were infected at an MOI = 0.01 with parental WT (WT 94-49), recombinant WT (rWT 94-49), or ΔSH (ΔSH 94-49) HMPV 94-49. Supernatant was removed every day out to 6 days and titered to measure extracellular virus. Error bars are +/-SD for technical duplicates.

**Figure S4.**
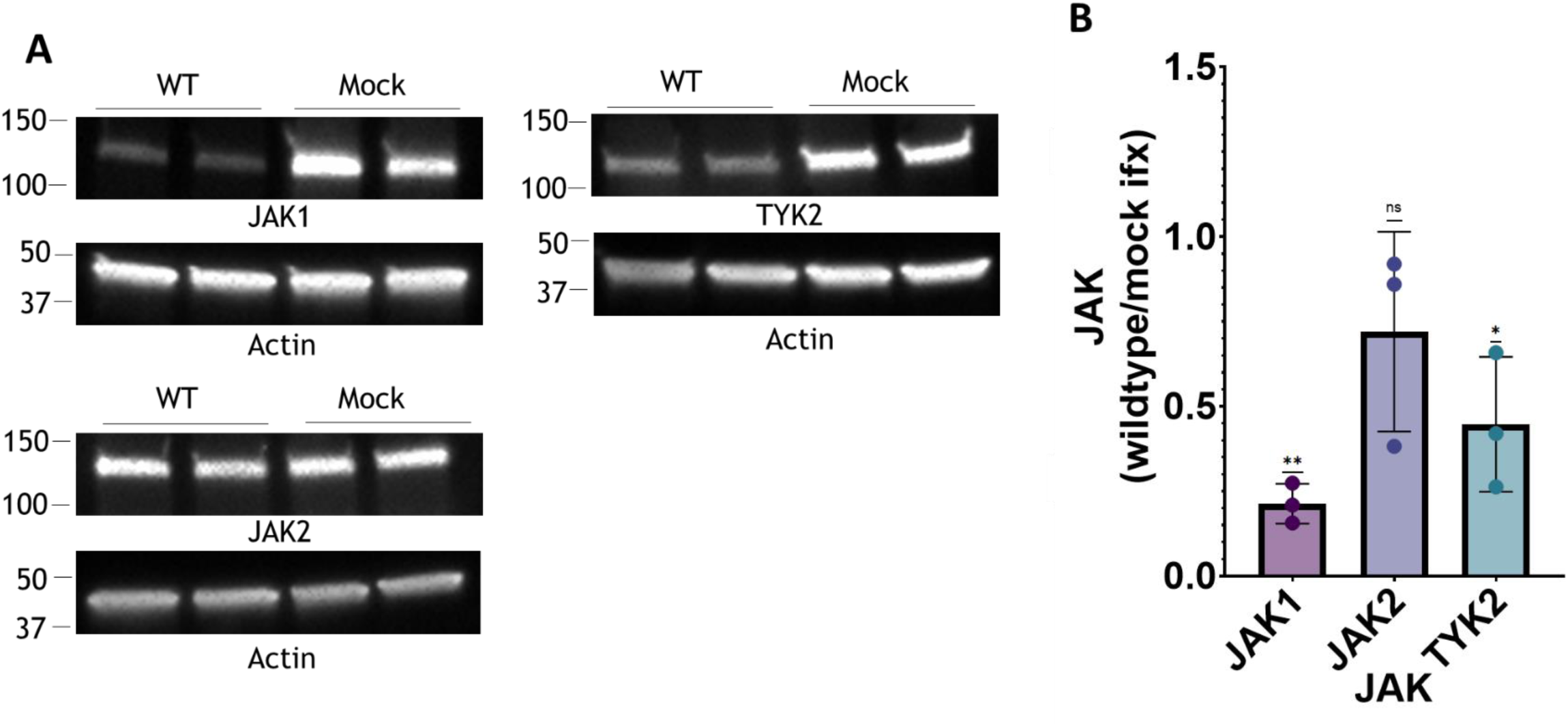
HMPV infection reduces JAK1 and TYK2. A549 cells were infected at an MOI of 0.5 for 24 hours & JAK levels were assessed by western blot. JAK1 levels in wildtype relative to mock infection conditions were assessed (B). Statistical analysis represents a one sample T-test (tested vs 1.0) performed on three biological replicates. Error bar represents SEM. * = P<0.05, **= P<0.01.

**Figure S5.**
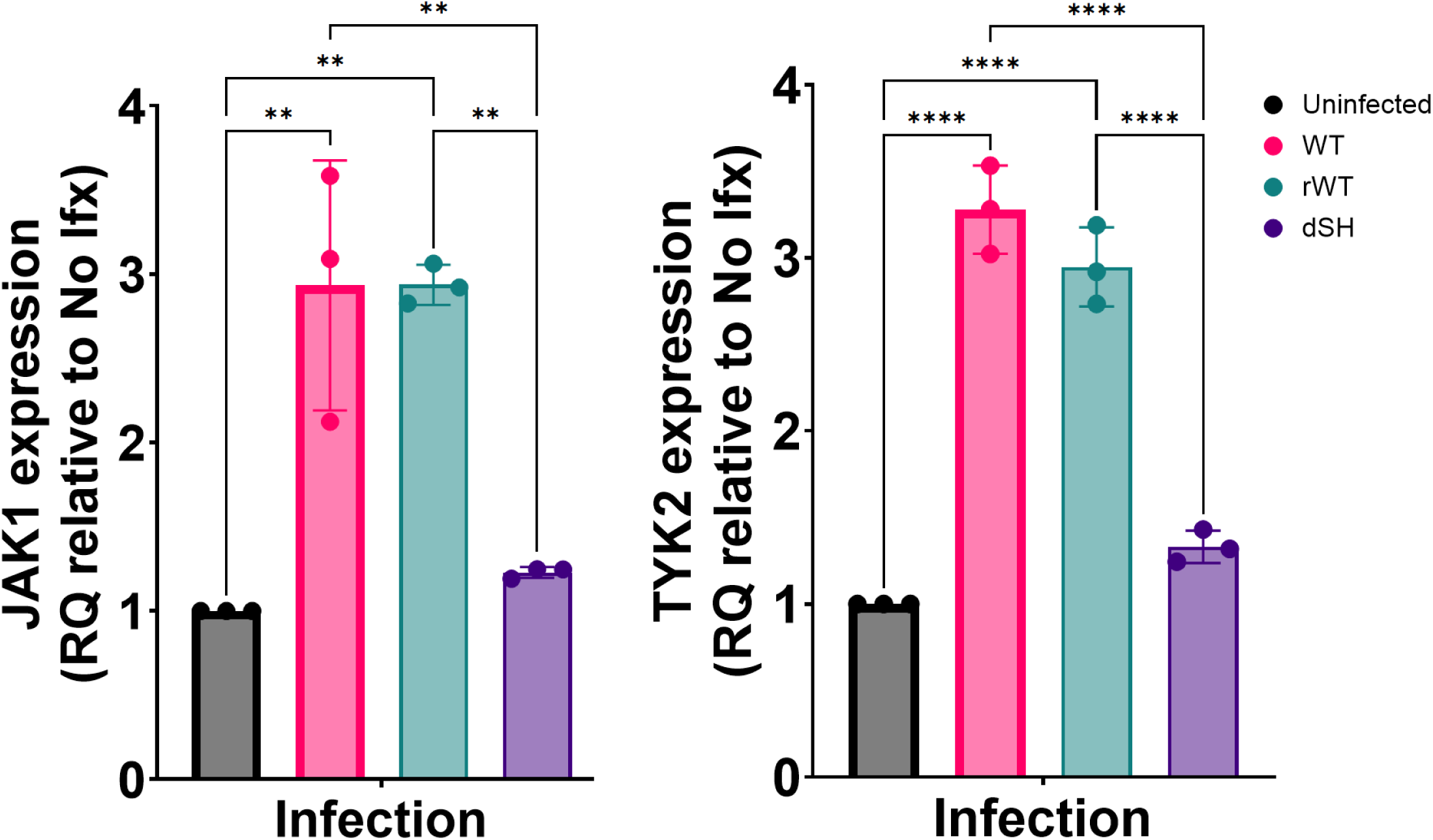
A549 cells were infected for 24h at MOI = 0.5 with parental wildtype 94-49 (WT), recombinant wildtype 94-49 (rWT), ΔSH 94-49 (ΔSH), or mock infection. JAK1 and TYK2 RNA levels were quantified via qPCR and normalized to mock infection. Error bars are +/-SD, with bars representing 3 biological replicates. Statistical comparisons are one-way ANOVAs. **= p<0.01, **** = p<0.0001. Non-significant comparisons hidden.

**Figure S6.**
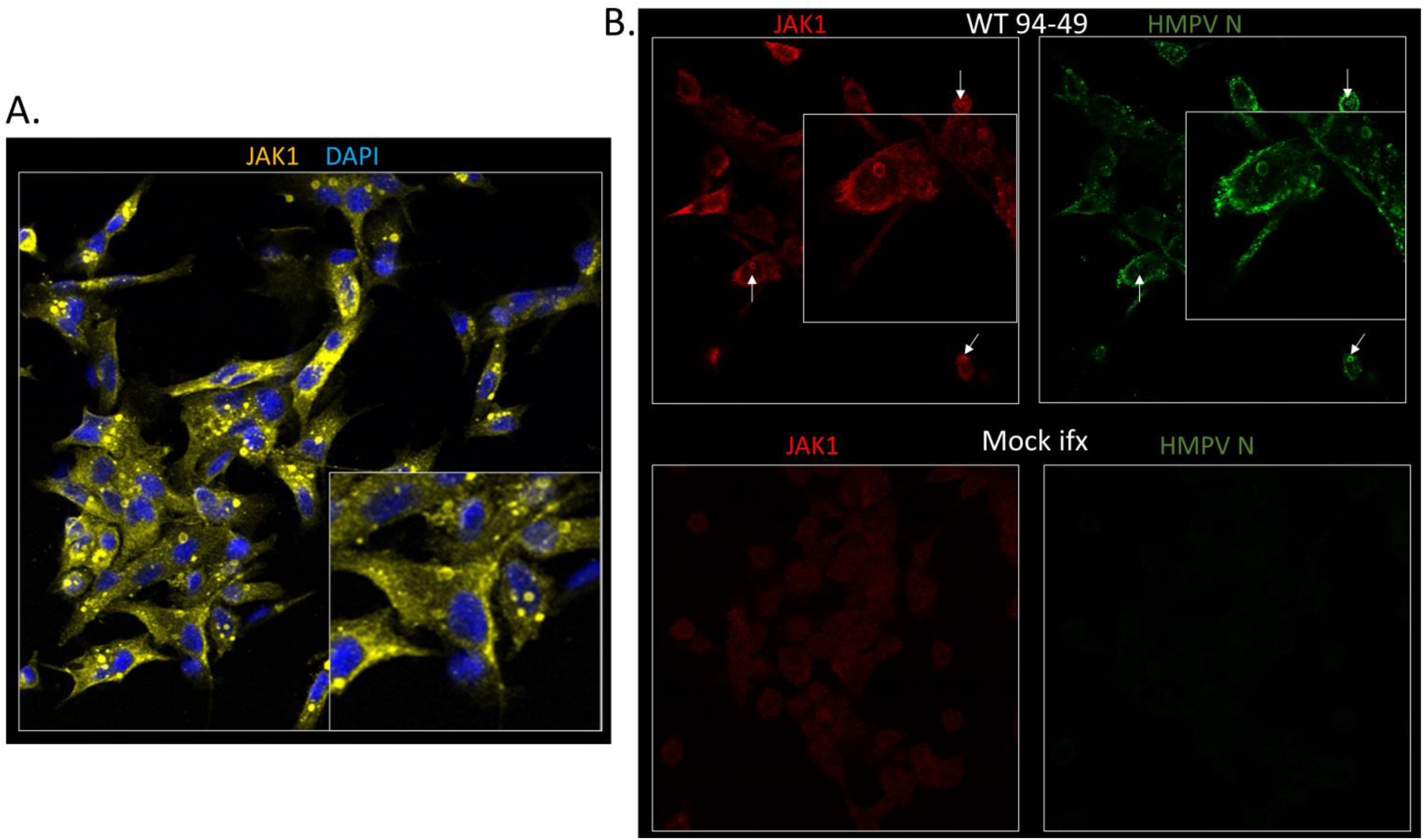
JAK1 localizes to viral inclusion bodies during infection. A549 cells were infected for 24 hours with an MOI of 0.5 of parental WT HMPV or mock (B) and stained for JAK1 (A) or JAK1 monoclonal (Millipore Sigma anti-JAK1 clone 73) with arrows added to show select colocalizing JAK1-HMPV N puncta (B). Enlarged insets added to show puncta details.

